# Biochemical reconstitution of branching microtubule nucleation

**DOI:** 10.1101/700047

**Authors:** Raymundo Alfaro-Aco, Akanksha Thawani, Sabine Petry

## Abstract

Microtubules are nucleated from specific locations at precise times in the cell cycle. However, the factors that constitute these microtubule nucleation pathways still need to be identified along with their mode of action. Here, using purified *Xenopus laevis* proteins we biochemically reconstitute branching microtubule nucleation, a nucleation pathway where microtubules originate from pre-existing microtubules, which is essential for spindle assembly and chromosome segregation. We found that besides the microtubule nucleator gamma-tubulin ring complex (γ-TuRC), the two branching effectors augmin and TPX2 are required to efficiently nucleate branched microtubules. Specifically, TPX2 generates regularly-spaced patches that recruit augmin and γ-TuRC to microtubules, which then nucleate new microtubules at preferred branching angles of less than 90 degrees. Our work demonstrates how γ-TuRC is brought to its nucleation site for branching microtubule nucleation. It provides a blueprint for other microtubule nucleation pathways and for generating a particular microtubule architecture by regulating microtubule nucleation.

## Introduction

Microtubules are nucleated from specific locations in the cell, and several of these microtubule nucleation pathways converge to form a particular cytoskeletal architecture (Kollman *et al.*, 2011; Lin *et al.*, 2015; Lüders and Stearns, 2007). Importantly, microtubules in cells are nucleated by the microtubule nucleator γ-TuRC (Kollman *et al.*, 2011; Zheng *et al.*, 1995) and its co-nucleation factor XMAP215 (Thawani *et al.*, 2018). At the same time, each microtubule nucleation pathway requires a unique set of nucleation effectors to recruit and regulate γ-TuRC at distinct cellular locations (Lin *et al.*, 2015). The identity of most of these effectors remains elusive, along with a mechanistic understanding of how they constitute the different microtubule nucleation pathways that generate the cytoskeleton.

Microtubules can nucleate from pre-existing microtubules, termed branching microtubule nucleation (Petry *et al.*, 2013), which amplifies microtubule number while preserving their polarity, as is needed in the mitotic spindle and in axons (Cunha-Ferreira *et al.*, 2018; David *et al.*, 2019; Kamasaki *et al.*, 2013; Petry *et al.*, 2013; Sánchez-Huertas *et al.*, 2016). The eight-subunit protein complex augmin is required for branching microtubule nucleation in plant, human and Drosophila cells, and meiotic Xenopus egg extract, where its depletion leads to reduced spindle microtubule density, less kinetochore fiber tension, metaphase arrest, and cytokinesis failure (David *et al.*, 2019; Decker *et al.*, 2018; Goshima *et al.*, 2008; Hayward *et al.*, 2014; Ho *et al.*, 2011; Kamasaki *et al.*, 2013; Lawo *et al.*, 2009; Nakaoka *et al.*, 2012; Petry *et al.*, 2011; Uehara *et al.*, 2009). Augmin is necessary to recruit γ-TuRC to spindle microtubules (Goshima *et al.*, 2007), and following the recombinant expression of augmin (Hsia *et al.*, 2014), this activity was confirmed using purified proteins (Song *et al.*, 2018). In meiotic Xenopus egg extract, the Ran-regulated protein TPX2 is released near chromatin (Gruss *et al.*, 2001), where it stimulates branching microtubule nucleation (Petry *et al.*, 2013), potentially by activating γ-TuRC (Alfaro-Aco *et al.*, 2017). Recently, TPX2 was also observed to form a co-condensate with tubulin along microtubules, which enhances the kinetic efficiency of branching microtubule nucleation (King and Petry, 2019). In meiotic Xenopus egg extract, TPX2 binds to microtubules before augmin/ γ-TuRC, followed by the nucleation event (Thawani *et al.*, 2019). In contrast, in mitotic Drosophila cells TPX2 is not required, and augmin binds to microtubules before γ-TuRC (Verma and Maresca, 2019). Despite numerous studies, exactly how augmin, TPX2 and γ-TuRC mediate branching microtubule nucleation, and whether they alone constitute a minimal system that nucleates branched microtubules, remains unclear. Here, we use biochemical reconstitution of its purified components to dissect branching microtubule nucleation mechanistically.

## Results and Discussion

Branching microtubule nucleation has been studied in Xenopus egg extract, where it is elicited by the constitutively active version of Ran (RanQ69L) (Petry *et al.*, 2013). In order to establish a controlled, minimal assay that furthers our mechanistic insight, we exposed a microtubule tethered to glass to sequential reaction mixtures of decreasing complexity and thereby regulated the availability of proteins necessary to stimulate branching microtubule nucleation. Using multicolor time-lapse total internal reflection (TIRF) microscopy, we first confirmed that an endogenous, pre-existing microtubule can serve as a template for branching microtubule nucleation when exposed to Ran-supplemented extract that releases branching factors (Figure 1A and Video 1). This shows that a microtubule formed independent of Ran can serve as the site for binding of branching factors and subsequent nucleation events. To gain mechanistic insight, we hypothesized that all necessary Ran-regulated branching factors bind to the pre-existing microtubule prior to microtubule nucleation. To test this, Ran-regulated branching factors were allowed to bind to taxol-stabilized pre-existing microtubules in the presence of nocodazole, which inhibits new microtubule formation (Figure 1B and Figure 1 – figure supplement 1). When another extract reaction was subsequently added, new microtubules nucleated almost exclusively from pre-existing microtubules, indicating that Ran-regulated branching factors bind to microtubules independent of their successful nucleation reactions (Figure 1B). Importantly, when RanQ69L was omitted and no branching factors were released in the second extract reaction, pre-existing microtubules simply elongated and branching microtubule nucleation was absent in the third reaction (Figure 1 – figure supplement 2A). To further test whether these microtubule-bound branching factors are sufficient for generating branches, we reduced the complexity of our assay by introducing only purified tubulin and GTP in the third reaction step. Surprisingly, short branched microtubules nucleated from pre-existing microtubules (Figure 1 – figure supplement 2B), showing that upon localization of branching factors and γ-TuRC, tubulin is the only protein required to form new branched microtubules from the localized factors. Further addition of XMAP215 to the final tubulin reaction made the short branches grow longer (Figure 1C). This revealed that branched microtubules retained the polarity of the pre-existing microtubule, and new microtubules do not appear to nucleate from other branched microtubules, suggesting that the branching factors do not relocate between microtubules (Figure 1C and Video 2). Thus, solely the deposition of branching factors and γ-TuRC to the pre-existing microtubule determines branching architecture.

**Figure 1.**
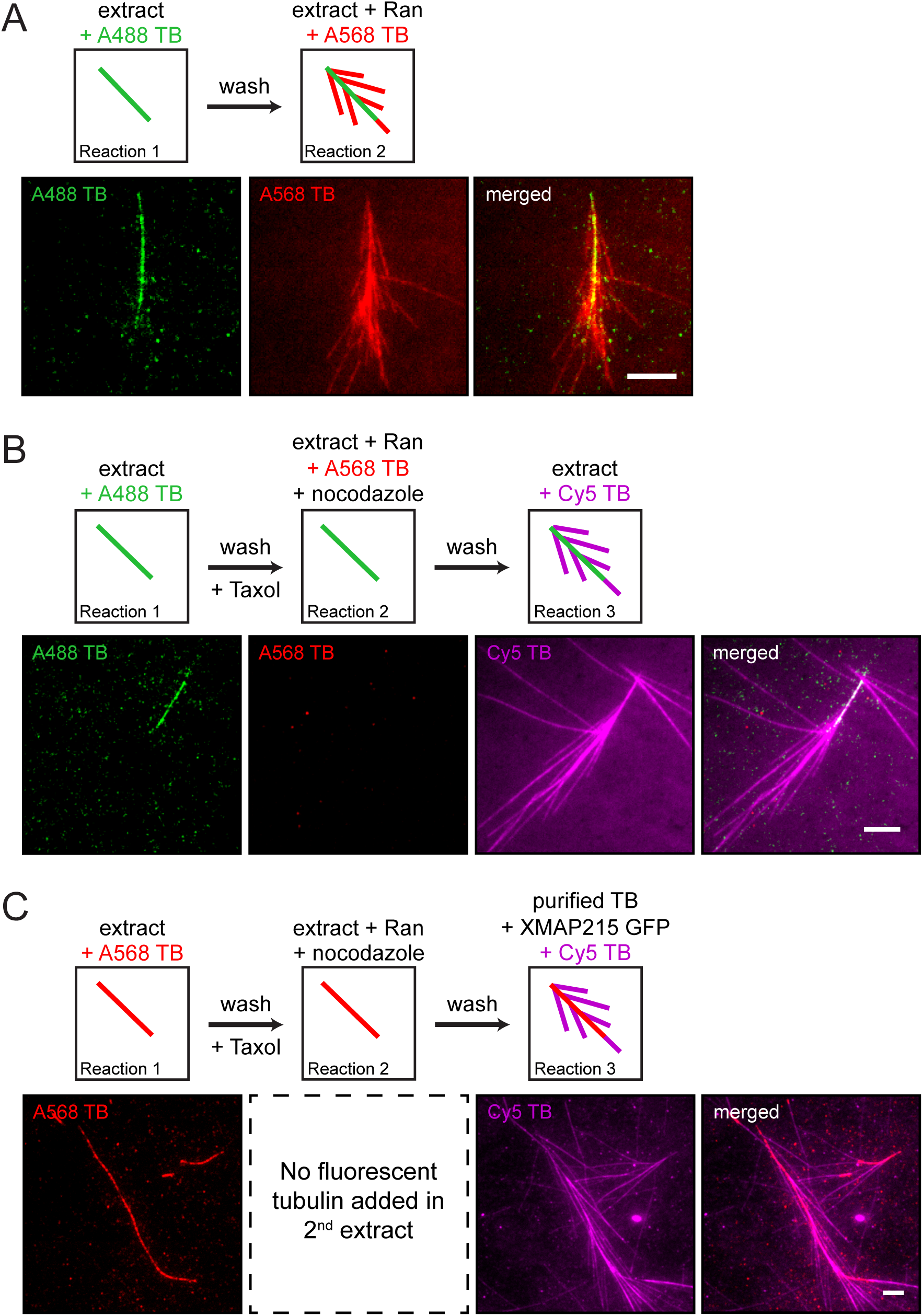
The proteins necessary for branching microtubule nucleation in Xenopus egg extract bind to a pre-existing microtubule independent of the nucleation event. (**A-C**) Sequential reactions with Xenopus egg extract. (**A**) Single microtubules formed on the glass surface in the first extract supplemented with Alexa488 tubulin (green). A second extract supplemented with Alexa568 tubulin (red) and RanQ69L was subsequently introduced. New microtubules (red) nucleated from pre-existing microtubules (green). See Video 1. (**B**) Single microtubules formed on the glass surface in the first extract supplemented with Alexa488 tubulin (green). A second extract supplemented with Alexa568 tubulin (red), RanQ69L and nocodazole was subsequently introduced, followed by a third extract supplemented with Cy5 tubulin (magenta). Branched microtubules (magenta) nucleated from pre-existing microtubules (green) via the branching factors released in the second extract, while no microtubules formed in the presence of nocodazole (red). See Figure 1 – figure supplement 1 and Figure 1 – figure supplement 2A. (**C**) Similar to (B), except that the first extract was supplemented with Alexa568 tubulin (red), the second extract contained no fluorescent tubulin, and the third extract reaction was substituted for purified Cy5 tubulin (magenta) and XMAP215. Branched microtubules (magenta) nucleated from pre-existing microtubules (red), which had been pre-loaded with branching factors in the second extract. See Figure 1 – figure supplement 2B and Video 2. For all experiments, images were collected approximately 5 min after the last solution was exchanged. Scale bars, 5 μm. The experiments were repeated three times with different Xenopus egg extracts.

Because the key for branching microtubule nucleation is to target γ-TuRC along the length of a microtubule, we tethered purified γ-TuRC along the microtubule lattice via artificial linkers, where it can still nucleate microtubules as a proof of concept (Figure 2 – figure supplement 1A-B). Therefore, if all branching factors are known, branching microtubule nucleation from a template microtubule can be reconstituted using purified components. To test this, we purified the essential proteins for branching microtubule nucleation in Xenopus egg extract (Petry *et al.*, 2013). The GFP-labeled eight-subunit *X. laevis* augmin holocomplex was co-expressed in insect cells and co-purified (Song *et al.*, 2018), the native 2.2 MDa γ-TuRC was purified from Xenopus egg extract (Thawani *et al.*, 2018), and GFP-TPX2 was expressed from *E. coli* and purified (King and Petry, 2019).

First, we assessed how the nucleator γ-TuRC gets targeted along the microtubule lattice. Purified TPX2, augmin and γ-TuRC in various combinations were added to surface-bound, GMPCPP-stabilized microtubules and imaged via TIRF microscopy (Figure 2A). γ-TuRC, visualized by a fluorescently-labeled antibody, bound along the length of microtubules in the presence of augmin (Figure 2 – figure supplement 2A-B) consistent with previous studies (Song *et al.*, 2018). Interestingly, more γ-TuRC was recruited along the microtubule lattice in the presence of both TPX2 and augmin (Figure 2B-C). Surprisingly, augmin and TPX2 formed distinct puncta on microtubules, where γ-TuRC was recruited. Using negative stain electron microscopy, we confirmed that γ-TuRC is recruited to regularly spaced patches, where it accumulates (Figure 2D). Next, we tested whether the microtubule binding proteins augmin and TPX2 need to bind in a certain sequence. Surprisingly, microtubule-bound TPX2 increased the amount of augmin bound to the microtubule, whereas the presence of augmin did not change the level of bound TPX2 (Figure 2E-F).

**Figure 2.**
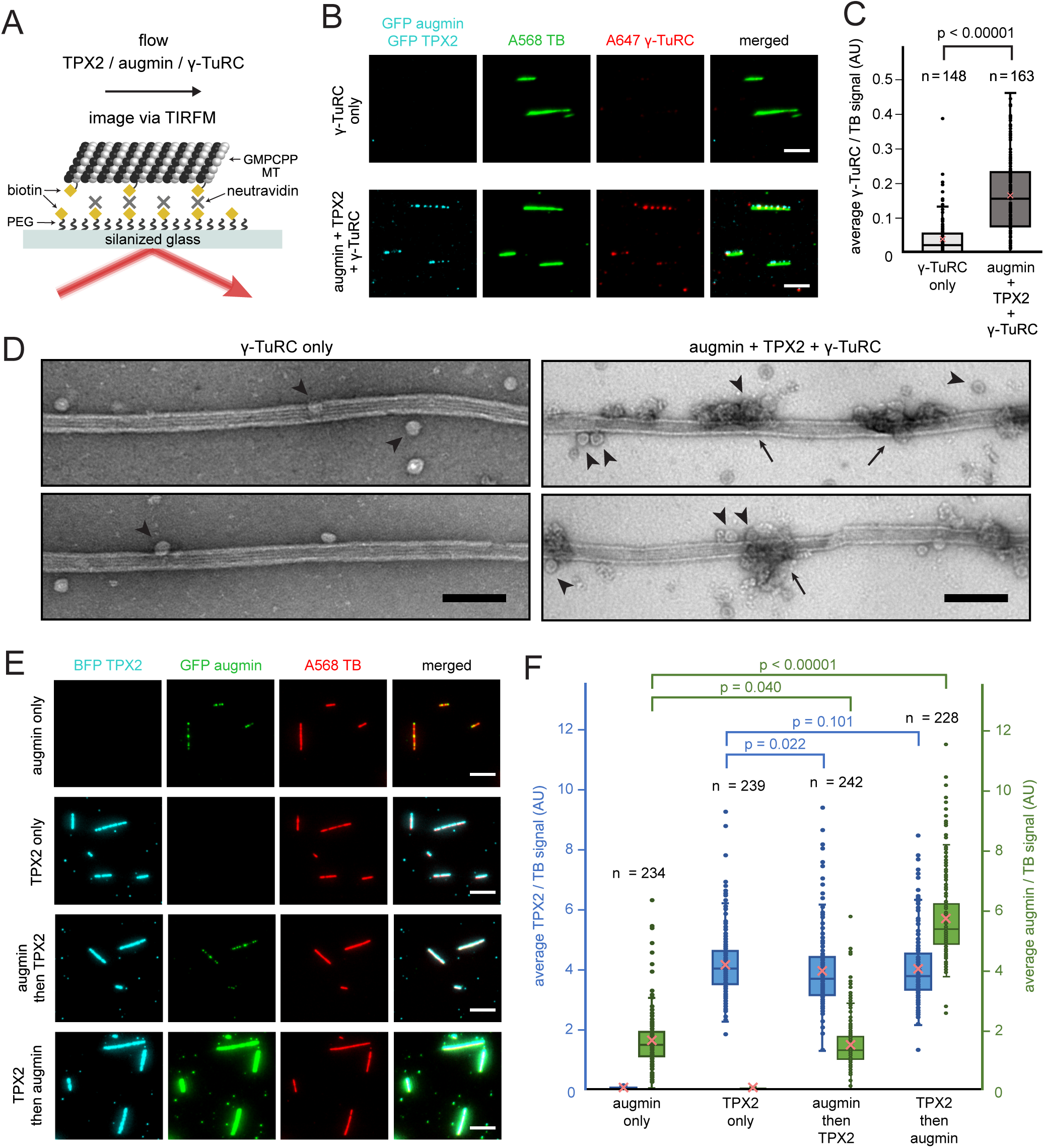
Binding of augmin, TPX2 and γ-TuRC to a template microtubule. (**A**) Diagram of the experimental set-up. GMPCPP-stabilized microtubules were attached to a PEG-passivated cover glass with biotin-neutravidin links. (**B**) γ-TuRC visualized using Alexa647-labeled antibodies (red) along microtubules (green), in the absence or presence of GFP-augmin and GFP-TPX2 (cyan). Scale bars, 5 μm. See Figure 2-figure supplement 2A-B. (**C**) Boxplot of average γ-TuRC signal relative to the average tubulin signal, where each dot represents one microtubule from the experiment in (B). The number of microtubules (n) was obtained from two replicates. (**D**) GMPCPP-stabilized microtubules incubated with γ-TuRC only or with augmin, TPX2 and γ-TuRC, visualized by electron microscopy after uranyl acetate staining. Ring-shaped structures that correspond to γ-TuRCs (arrowheads), and clusters of protein formed on microtubules (arrows) are visible. Scale bars, 100 nm. (**E**) GFP-augmin (green) and BFP-TPX2 (cyan) visualized along microtubules (red) by themselves or in sequential binding steps. Scale bars, 5 μm. (**F**) Boxplot of average BFP-TPX2 signal or GFP-augmin signal relative to the average tubulin signal, where each dot represents one microtubule from the experiment in (E). The number of microtubules (n) was obtained from two replicates. For (C) and (F), the boxes extend from 25th to 75th percentiles, the whiskers extend from minimum to maximum values, and the mean values are plotted as crosses. P-values were calculated from independent T-tests.

Having established that purified TPX2 and augmin recruit γ-TuRC to template microtubules, can they indeed cause branching microtubule nucleation? All three factors were bound to a stabilized microtubule as above, followed by addition of tubulin and GTP in polymerization buffer (Figure 3A). Remarkably, branching microtubule nucleation from a template microtubule occurred using only purified proteins (Figure 3B and Video 3). Live microscopy allowed us to accurately distinguish branching microtubule nucleation from microtubules that were spontaneously nucleated before contacting the microtubule template (Figure 3 – figure supplement 1A). Thus, TPX2, augmin and γ-TuRC are sufficient to specifically nucleate new branched microtubules, which remain attached at the nucleation site on the template microtubule (Figure 3B). In rare instances, enough microtubules branch from a single microtubule template to create structures reminiscent of those from Xenopus egg extract (Figure 3B, bottom).

**Figure 3.**
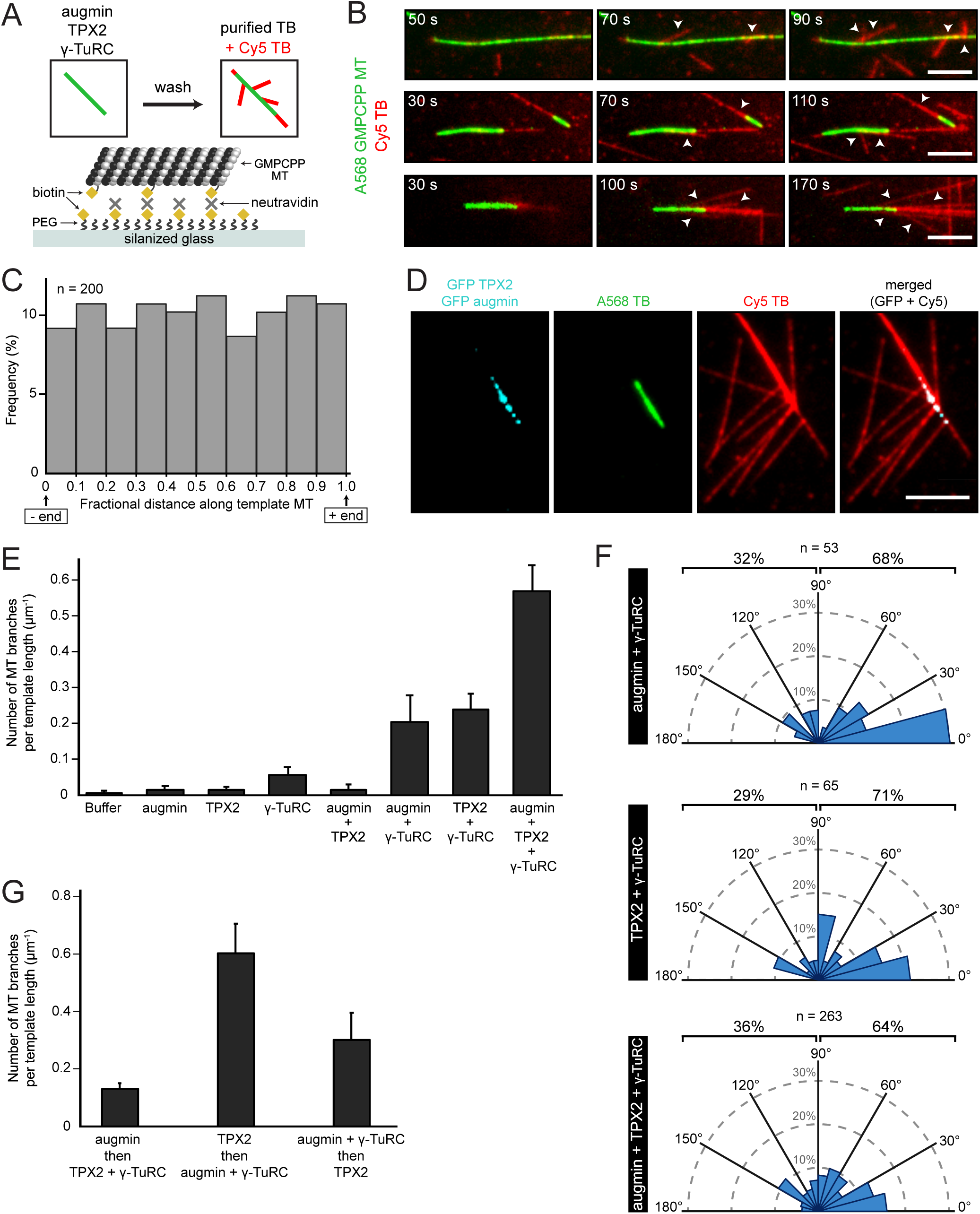
Biochemical reconstitution of branching microtubule nucleation using purified augmin, TPX2 and γ-TuRC. (**A**) Diagram of the experimental set-up. GMPCPP-stabilized microtubules were attached to a PEG-passivated cover glass with biotin-neutravidin links. Following the binding of augmin, TPX2, and γ-TuRC, nucleation of new microtubules was visualized using Cy5 tubulin. (**B**) Using the set-up in (A), the formation of microtubule branches (red, arrowheads) from GMPCPP-stabilized microtubules (green) was observed. Scale bars, 5 μm. See Figure 3 – figure supplement 1A and Video 3. (**C**) Fractional distance along the template microtubule where microtubule branches formed. The 0-point on the x-axis denotes nucleation at the minus-end of the template microtubule, while the 1-point denotes nucleation at the plus-end. The number of branching events (n) was obtained from twelve replicates using γ-TuRC purified from four different preps. (**D**) Same as (A), microtubule branches (red) grow from distinct GFP-augmin and GFP-TPX2 puncta (cyan) localized on GMPCPP-stabilized microtubules (green). (**E**) Number of microtubule branches per field of view after 4 min, normalized to the length of template microtubule available, for all the combinations of branching factors. Values are the mean of four replicates using γ-TuRC purified from one prep, and error bars represent standard error of the mean. See Figure 3 – figure supplement 1B-C. (**F**) Angle of branching for three different combinations of branching factors. The number of branching events (n) was obtained from eight replicates using γ-TuRC purified from two different preps in the case of augmin + γ-TuRC and TPX2 + γ-TuRC, and from twelve replicates using γ-TuRC purified from four different preps in the case of augmin + TPX2 + γ-TuRC. (**G**) Number of microtubule branches per field of view after 4 min, normalized to the length of template microtubule available, for different binding sequences. Values are the mean of four replicates using γ-TuRC purified from one prep, and error bars represent standard error of the mean.

How are the nucleation sites spatially organized along the template microtubule? New microtubules nucleate all along the template microtubule without any preference for the template’s plus- or minus-ends (Figure 3C), likely because the template microtubule was fully available for simultaneous binding of branching factors in our assay set-up. Thus, there is no signature on the stabilized microtubule lattice that determines where a branch occurs, only that microtubule nucleation events occur from distinct TPX2/augmin puncta distributed along the microtubule lattice (Figure 3D). Multiple microtubules can be generated from the same puncta as resolved by light microscopy (Figure 3D), which presumably nucleated from neighboring γ-TuRCs (Figure 2D).

What does each protein contribute to branching microtubule nucleation? To test this, each purified factor was assessed alone for its nucleation potential from a template microtubule, combined in pairs and ultimately altogether. Notably, γ-TuRC is essential for branching microtubule nucleation (Figure 3E). Despite the fact that TPX2 can recruit tubulin (King and Petry, 2019), it alone or together with augmin cannot nucleate branched microtubules. γ-TuRC can infrequently bind to the microtubule lattice on its own (Figure 2D), leading to rare nucleation events without TPX2 and augmin (Figure 3E). Not surprisingly, augmin and γ-TuRC can cause branching microtubule nucleation to a limited extent (Figure 3E and Figure 3 – figure supplement 1B), as augmin can directly recruit γ-TuRC to a pre-existing microtubule *in vitro* (Song *et al.*, 2018). Surprisingly, TPX2 and γ-TuRC can also cause branching microtubule nucleation to a similar extent as augmin and γ-TuRC (Figure 3E and Figure 3 – figure supplement 1C). Importantly, only when augmin, TPX2 and γ-TuRC are present, branching microtubule nucleation occurs most often (Figure 3E). Branched microtubules are preferentially formed in angles < 90 degrees, with 0-15 degrees being the most common (Figure 3F). This way, most branched microtubules maintain the same polarity as the mother microtubule, a hallmark of branching microtubule nucleation. Interestingly, the angle of microtubule branches did not drastically change when augmin or TPX2 were combined with γ-TuRC, only that augmin/γ-TuRC alone caused a higher proportion of shallow branch angles (Figure 3F).

Next, we tested whether branching microtubule nucleation is further enhanced by having XMAP215 present. Indeed, XMAP215 co-localizes to the template microtubule and appears to increase both microtubule nucleation rate and growth (Figure 3 – figure supplement 2). Exact quantification of this effect was not possible because branched microtubules were already formed before imaging was possible and microtubules quickly grew into each other, preventing the accurate identification of branching microtubule nucleation. Lastly, knowing that the binding sequence of TPX2 and augmin matters for maximum factor recruitment, does this have an effect on nucleation? Indeed, only when TPX2 was bound first and augmin/γ-TuRC second, a higher level of branching microtubule nucleation was measured (Figure 3G).

Via an *in vitro* reconstitution, we demonstrate that the three factors TPX2, augmin and γ-TuRC are sufficient to cause branching microtubule nucleation and defined the roles of each protein. Interestingly, augmin and γ-TuRC alone can nucleate branched microtubules. This may be reflective of cell types where TPX2 is not needed for branching microtubule nucleation, such as mitotic Drosophila cells (Verma and Maresca, 2019). Unexpectedly, it is beneficial that TPX2 binds to microtubules first, as it recruits more augmin, in addition to its ability to recruit tubulin (King and Petry, 2019). This is similar to a recent observation in Xenopus egg extract where TPX2 binds to microtubules first, and augmin cannot bind to microtubules in its absence (Thawani *et al.*, 2019). This implies that TPX2 directly regulates augmin’s binding to microtubules, but does not rule out additional regulation that could encompass other factors such as EML3 (Luo *et al.*, 2019).

Although microtubule nucleation effectors alone can generate microtubules *in vitro* (Roostalu *et al.*, 2015; Woodruff *et al.*, 2017), γ-TuRC is required for physiological microtubule nucleation (Kollman *et al.*, 2011; Thawani *et al.*, 2018), and this reconstitution highlights the importance of including it when studying microtubule nucleation. Localizing γ-TuRC to a specific location, from which it nucleates a microtubule *in vitro* as it occurs in the cell, serves as a pioneering example for the biochemical reconstitution of other microtubule nucleation pathways or microtubule organizing centers. It is analogous to the *in vitro* reconstitution of actin branching (Mullins *et al.*, 1998), which paved the way to explain how the actin cytoskeleton supports cell function. This work serves as a platform to study how microtubule nucleation creates different microtubule architectures and will allow reconstituting larger structures based on this microtubule nucleation pathway, such as the mitotic spindle.

## Materials and Methods

### Cloning, expression and purification of proteins

DH5α *E. coli* cells (New England Biolabs, C2987I) were used for all cloning steps. Rosseta2 (DE3)pLysS cells (Novagen, 714034) were used for all protein expression in *E. coli*, and cultures were grown in TB Broth (Sigma-Aldrich, T0918), or in LB Broth (Sigma-Aldrich, L3522) for the expression of TPX2. Sf9 cells using the Bac-to-Bac system (Invitrogen) were used in the expression of augmin and XMAP215, and cultures were grown in Sf-900 III SFM (Gibco, 12658027).

Human RanQ69L with N-terminal Strep-6xHis-BFP, and human EB1 with C-terminal GFP-6xHis were expressed and purified as previously described (Thawani *et al.*, 2019). Full-length *Xenopus laevis* TPX2 constructs were expressed and purified as previously described (King and Petry, 2019). Briefly, N-terminal Strep-6xHis-GFP and Strep-6xHis-BFP were cloned into pST50 vectors and expressed in *E. coli* for 7 hr at 25°C. Both proteins were affinity purified using Ni-NTA agarose beads (Qiagen, 30250) followed by gel filtration with a Superdex 200 HiLoad 16/600 column (GE Healthcare) in CSF-XB buffer (100 mM KCl, 10 mM K-HEPES, 1 mM MgCl_2_, 0.1 mM CaCl_2_, 5 mM EGTA, pH 7.7) + 10% w/v sucrose. Full-length *Xenopus laevis* XMAP215 with C-terminal GFP-7xHis was expressed in Sf9 cells using the Bac-to-Bac system and purified as previously described (Thawani *et al.*, 2018). Briefly, XMAP215 was affinity-purified using a HisTrap HP 5 ml column (GE Healthcare), followed by cation-exchange with a Mono S 10/100 GL column (GE Healthcare). The protein was dialyzed overnight into CSF-XB + 10% w/v sucrose. GFP-tagged *Xenopus laevis* augmin holocomplex was co-expressed in Sf9 cells using the Bac-to-Bac system and purified as previously described (Song *et al.*, 2018). Briefly 1–2 liters of Sf9 cells (1.5–2.0 × 10^6^ mL^-1^) were co-infected with different baculoviruses, each carrying a subunit of the augmin complex, at MOIs of 1–3. Cells were collected 72 h after infection. HAUS6 had an N-terminal ZZ-tag and HAUS2 had a C-terminal GFP-6xHis. The remaining six subunits were untagged. Augmin holocomplex was affinity-purified using IgG-Sepharose (GE Healthcare, 17-0969-01) and eluted via cleavage with 100–200 µg of GST-HRV3C protease. The HRV3C protease was subsequently removed using a GSTrap 5mL column (GE Healthcare). The sample was further purified and concentrated using Ni-NTA agarose beads. The protein was dialyzed overnight into CSF-XB + 10% w/v sucrose. All recombinant proteins were flash frozen and stored at −80 °C. Protein concentrations were determined with Bradford dye (Bio-Rad, 5000205) or using a Coomassie-stained SDS-PAGE gel loaded with known concentrations of BSA (Sigma-Aldrich, B6917).

Native γ-TuRC was purified from Xenopus egg extract with some changes to previously described protocols (Zheng *et al.*, 1995; Thawani *et al.*, 2018). 5 ml of Xenopus egg extract were diluted 10-fold with CSF-XB + 10% w/v sucrose, 1 mM GTP, 1 mM DTT, and 10 μg ml^-1^ leupeptin, pepstatin and chymostatin. Large particles were removed by spinning at 3000 *g* for 10 min at 4°C. The supernatant was further diluted two-fold with buffer and passed through filters of decreasing pore size (1.2 µm, 0.8 µm and 0.22 µm). γ-TuRC was precipitated from the filtered extract by addition of 6.5% w/v polyethylene glycol (PEG) 8000 and incubated on ice for 30 min. After centrifugation for 20 min at 17,000 *g* at 4°C, the pellet was resuspended in 15 ml of the initial CSF-XB buffer supplemented with 0.05% NP-40. The resuspended pellet was centrifuged at 136,000 *g* at 4°C for 7 min. The supernatant was then precleared with protein A Sepharose beads (GE Healthcare, 45002971) for 20 min at 4°C. The beads were removed by spinning, 2-4 ml γ-tubulin antibody (1 mg ml^-1^) was added to the sample, and the sample was rotated at 4°C for 2 h. After this, 1 ml of washed Protein A Sepharose beads was incubated with the sample on the rotator for 2 h at 4°C. The beads were collected by spinning, and subsequently transferred to a column with the same buffer used to resuspend the PEG pellet. The beads were washed with the initial CSF-XB buffer supplemented with extra 150 mM KCl, then with CSF-XB buffer supplemented with 1 mM ATP, and finally with CSF-XB buffer to remove the ATP. For biotinylation of γ-TuRC, the beads were incubated with 25 µM of NHS-PEG4-biotin (Thermo Scientific, A39259) in CSF-XB buffer for 1 h at 4°C, and unreacted reagent was washed away with CSF-XB buffer before elution with γ-tubulin peptide. 2 ml γ-tubulin peptide (amino acids 412–451) at 0.5 mg ml^-1^ in CSF-XB buffer was applied to the column and allowed to incubate overnight. The eluted sample was collected the following day, and it was concentrated using a 100 kDa MWCO centrifugal-filter (Amicon, UFC810024). This concentrated sample was loaded onto a 10-50% w/w sucrose gradient in the initial CSF-XB buffer, and centrifuged at 200,000 *g* for 3 h at 4°C in a TLS55 rotor (Beckman Coulter). The sucrose gradient was fractionated manually from the top, and the fractions with the highest γ-tubulin signal by Western blotting were combined and concentrated using another 100 kDa MWCO centrifugal-filter. Purified γ-TuRC was always used within two days on ice without freezing.

Unlabeled cycled tubulin purified from bovine brain was obtained from a commercial source (PurSolutions, 032005). Before use, all proteins were pre-cleared of aggregates via centrifugation at 80,000 RPM for 15 min at 4°C in a TLA100 rotor (Beckman Coulter).

### Tubulin labeling and polymerization of GMPCPP-stabilized microtubules

Bovine brain tubulin was labeled following previously described methods (Hyman *et al.*, 1991). Using Cy5-NHS ester (GE Healthcare, PA15101) yielded 54-70% labeling. Using Alexa-568 NHS ester (Invitrogen, A20003) yielded 36-40% labeling. Labeling efficiency with biotin-PEG4-NHS (Thermo Scientific, A39259) was not calculated.

Single-cycled GMPCPP-stabilized microtubules were made as previously described (Gell *et al.*, 2010). Briefly, 12 μM unlabeled tubulin + 1 μM Alexa-568 tubulin + 1 μM biotin tubulin was polymerized in BRB80 (80 mM Pipes, 1 mM EGTA, 1 mM MgCl_2_) in the presence of 1 mM GMPCPP (Jena Bioscience, NU-405L) for 1 h at 37°C. For GMPCPP-stabilized microtubules without any labels, 14 μM unlabeled tubulin was polymerized. For GMPCPP-stabilized microtubules without biotin, 13 μM unlabeled tubulin + 1 μM Alexa-568 tubulin was polymerized.

### Preparation of polyethylene glycol (PEG)-functionalized surfaces

Cover glasses (Carl Zeiss, 474030-9020-000) were silanized and reacted with PEG as previously described (Bieling *et al.*, 2010), except that hydroxyl-PEG-3000-amine (Rapp Polymere, 103000-20) and biotin-PEG-3000-amine (Rapp Polymere, 133000-25-20) were used. Glass slides were passivated with poly(L-lysine)-PEG (SuSoS) (Bieling *et al.*, 2010). Flow chambers for TIRF microscopy were assembled using double-sided tape.

### Attachment of GMPCPP-stabilized microtubules to PEG-functionalized surfaces

The assay was performed following a previously described protocol with some changes (Roostalu *et al.*, 2015). Flow chambers were incubated with 5% Pluronic F-127 in water (Invitrogen, P6866) for 10 min at room temperature and then washed with assay buffer (80 mM Pipes, 30 mM KCl, 1 mM EGTA, 1 mM MgCl_2_, 1 mM GTP, 5 mM 2-mercaptoethanol, 0.075% (w/v) methylcellulose (4,000 cP; Sigma-Aldrich, M0512), 1% (w/v) glucose, 0.02% (v/v) Brij-35 (Thermo Scientific, 20150)) supplemented with 50 μg mL^-1^ k-casein (Sigma-Aldrich, C0406) and extra 0.012% (v/v) Brij-35. Flow chambers were then incubated with assay buffer containing 50 μg mL^-1^ NeutrAvidin (Invitrogen, A2666) for 3 min on a metal block on ice and subsequently washed with BRB80 (80 mM Pipes, 1 mM EGTA, 1 mM MgCl_2_). Next, flow chambers were incubated for 5 min at room temperature with biotin- and Alexa-568-labeled GMPCPP-stabilized microtubules diluted 1:2000 in BRB80. Unbound microtubules were removed by subsequent BRB80 washes.

### Binding of proteins to GMPCPP-stabilized microtubules

To test the recruitment of γ-TuRC to a microtubule by augmin and TPX2, a mixture of GFP-TPX2 (50 nM), GFP-augmin (50 nM) and γ-TuRC, which was previously incubated for 5 min on ice, was added to a flow chamber that had GMPCPP-stabilized microtubules attached to the surface as described above. This was incubated for 5 min at room temperature. Unbound proteins were removed with additional BRB80 washes. To visualize native γ-TuRC, Alexa-647 (Invitrogen) labeled antibodies against γ-tubulin (XenC antibody, 2 μg ml^-1^) were added to the flow chamber and incubated for 10 min at room temperature. Unbound antibody was removed with additional BRB80 washes, and the final solution was exchanged to BRB80 + 250 nM glucose oxidase (Crescent Chemical, SE22778.02), 64 nM catalase (Sigma-Aldrich, C40) and 1% (w/v) glucose. The sample was imaged immediately. For experiments where one or two of the proteins in the mixture were omitted, the volume was substituted with CSF-XB buffer. The same set-up was used when imaging the binding of GFP-augmin (50 nM) and GFP-TPX2 (50 nM) to microtubules in the presence of each other. In these cases, γ-TuRC was not included, and instead of adding XenC antibody, BRB80 + oxygen scavengers were added after the last unbound proteins were removed by BRB80 washes. In experiments where two proteins were bound to microtubules sequentially, unbound protein was removed by BRB80 washes before the second protein was added.

### Microtubule nucleation assays on PEG-functionalized surfaces

For branching microtubule nucleation reactions *in vitro*, a mixture of TPX2 (50 nM), augmin (50 nM) and γ-TuRC, which was previously incubated for 5 min on ice, was added to the chamber containing attached GMPCPP-stabilized microtubules and incubated for 5 min at room temperature. Unbound proteins were removed by additional BRB80 washes. The final assay mixture was flowed into the chambers: 80 mM Pipes, 30 mM KCl, 1 mM EGTA, 1 mM MgCl_2_, 1 mM GTP, 5 mM 2-mercaptoethanol, 0.075% (w/v) methylcellulose (4,000 cP), 1% (w/v) glucose, 0.02% (v/v) Brij-35, 250 nM glucose oxidase, 64 nM catalase, 1 mg ml^-1^ BSA, 19 μM unlabeled bovine tubulin and 1 μM Cy5-labeled bovine tubulin. For experiments where XMAP215-GFP was added to this final reaction its concentration was 50 nM.

### Microtubule nucleation from artificially-attached γ-TuRCs to microtubules

Coverslips were coated with dichlorodimethylsilane (Gell *et al.*, 2010). Flow chambers for TIRF microscopy were assembled using double-sided tape and incubated for 5 min at room temperature with biotin- and Alexa-568-labeled GMPCPP-stabilized microtubules diluted 1:2000 in BRB80. A small number of microtubules attached non-specifically to the glass, and the rest were removed with BRB80 washes. The rest of the glass surface was blocked with 1% Pluronic F127, and the chamber was incubated for 3 min at room temperature with 500 μg mL^-1^ NeutrAvidin diluted in BRB80. Undiluted biotinylated γ-TuRC was incubated in the chamber for 10 min at room temperature, and after washing with BRB80 the final tubulin nucleation mix was added: 80 mM Pipes, 1 mM EGTA, 1 mM MgCl_2_, 1 mM GTP, 2.5 mM PCA, 25 nM PCD, 2 mM Trolox, 19 μM unlabeled bovine tubulin and 1 μM Cy5-labeled bovine tubulin.

### Sequential Xenopus egg extract reactions

CSF extracts were prepared from *Xenopus laevis* oocytes as described previously (Murray and Kirschner, 1989; Hannak and Heald, 2006). When working with *Xenopus laevis*, all relevant ethical regulations were followed, and all procedures were approved by Princeton IACUC. Extract reactions were done in flow chambers prepared between glass slides and 22 × 22 mm, 1.5 coverslips (Fisherbrand, 12-541B) using double-sided tape. In all reactions 75% of the total volume was extract, and 25% was a combination of other components or CSF-XB (100 mM KCl, 10 mM K-HEPES, 1 mM MgCl_2_, 0.1 mM CaCl_2_, 5 mM EGTA, pH 7.7) + 10% w/v sucrose. All reactions were done in the presence of 0.5 mM sodium orthovanadate (NEB, P0758S) to avoid sliding of microtubules on the glass surface, and with 0.89 µM fluorescently-labeled tubulin. In reactions where BFP-RanQ69L was added, its concentration was 10 µM. When EB1-GFP was added its concentration was 85 nM. All proteins and chemicals added to egg extracts were stored or diluted into CSF-XB buffer + 10% w/v sucrose. Reaction mixtures were pipetted into the flow chambers to initiate microtubule formation.

For sequential extract reactions, individual microtubules were allowed to form on the glass surface from the first extract reaction for 5-8 min, and soluble, non-microtubule bound proteins were removed by washing with CSF-XB. For experiments with three sequential reactions, the CSF-XB wash was supplemented with 0.05 mM Taxol (Sigma-Aldrich, T7402). The second extract reaction was then introduced. In the case of Fig. 1a the chamber was imaged immediately. For all other experiments, the second extract with 0.033 mM nocodazole (Sigma-Aldrich, M1404) was incubated in the chamber for 5 min, followed by the removal of unbound protein with CSF-XB if the third reaction was extract, or with BRB80 if the third reaction was purified tubulin. The third extract reaction was then introduced and imaged immediately. For experiments where the final reaction was purified tubulin, the final tubulin nucleation mix was added: 80 mM Pipes, 1 mM EGTA, 1 mM MgCl_2_, 1 mM GTP, 2.5 mM PCA, 25 nM PCD, 2 mM Trolox, 19 μM unlabeled bovine tubulin and 1 μM Cy5-labeled bovine tubulin. If XMAP215-GFP was added in this final reaction, its concentration was 25 nM.

### TIRF microscopy and image analysis

Total internal reflection fluorescence (TIRF) microscopy was performed with a Nikon TiE microscope using a 100x 1.49 NA objective. Andor Zyla sCMOS camera was used for acquisition, with a field of view of 165.1 × 139.3 µm. 2 × 2 binned, multi-color images were acquired using NIS-Elements software (Nikon). All adjustable imaging parameters (exposure time, laser intensity, and TIRF angle) were kept the same within experiments. For microtubule nucleation assays *in vitro* the TIRF objective was warmed to 33°C using an objective heater (Bioptechs, 150819-13). For all time-lapse imaging, multi-color images were collected every 2 seconds. Brightness and contrast were optimized individually for display, except for images in Fig. 2, Extended Data Fig. 4 and Extended Data Fig. 7, where images belonging to the same experiment were contrast-matched.

Images used for the quantification of microtubule binding were analyzed using ImageJ (Schindelin *et al.*, 2012). To segment microtubules, the tubulin signal was first thresholded via the Otsu method. Microtubules were isolated from the mask by setting the minimum particle area as 1 μm^2^. Average fluorescent signals per pixel, for the microtubule or bound proteins, were calculated for each microtubule. The average intensity from the reverse mask of the entire field of view was subtracted from the average intensity on each microtubule. For branching microtubule nucleation experiments *in vitro*, individual branching events were counted manually using time-lapse experiments within the first 3.5 min of the reaction. Lengths of microtubules and branching angles were measured using ImageJ.

### Negative stain electron microscopy

Unlabeled GMPCPP-stabilized microtubules diluted 1:500 were incubated for 5 min at room temperature with either γ-TuRC only or with a mixture of TPX2 (50 nM) + augmin (50 nM) + γ-TuRC. The samples were diluted 10-fold with BRB80 to reduce the number of unbound γ-TuRC molecules in the background, and 5 µl of this diluted sample was immediately applied onto glow-discharged grids (Electron Microscopy Sciences, CF400-Cu). The samples were stained with 2% uranyl acetate. Images were collected with a CM100 TEM (Philips) at 80 keV at a magnification of 64,000. Images were recorded using an ORCA camera.

### Antibodies

Polyclonal XenC antibody was a gift from C. Wiese and was described previously (Wiese and Zheng, 2000). It was used to generate Alexa-647-labeled XenC antibody by first dialyzing antibodies in PBS buffer (50 mM NaPO_4_, 150 mM NaCl, pH 7.4). The reaction with Alexa-647-NHS-ester was done according to the protocol recommended by the manufacturer. Finally, the removal of unreacted dye was done via gel filtration in Bio-Gel P-30 Gel (Bio-Rad). On average, each XenC antibody was labeled with 2.5 Alexa-647 dye molecules. The polyclonal antibody used to purify γ-TuRC from Xenopus egg extract was generated against a purified γ-tubulin peptide (amino acids 412-451) through a commercial vendor (Genscript). The presence of γ-TuRC during its purification was tracked via Western blotting using the GTU88 (Sigma-Aldrich, T6557) antibody against γ-tubulin.

## Acknowledgements

We are grateful to Christiane Wiese for providing XenC antibodies, Jae-Geun Song and Brian Mahon for help in the expression and purification of augmin, Matt King for help in the expression and purification of TPX2, all members of the Petry laboratory for discussions, Thomas Surrey for sharing the detailed protocol for making biotin-PEG-functionalized coverslips, and James Wakefield for sharing unpublished data and critically reading this manuscript. We thank Ron Vale with whom the original project vision was conceived. This work was supported by the HHMI Gilliam Fellowship and the NSF Graduate Research Fellowship (to R.A.), the American Heart Association predoctoral fellowship 17PRE33660328 (to A.T.), the NIH New Innovator Award, the Pew Scholars Program in the Biomedical Sciences, and the David and Lucile Packard Foundation (to S.P.).

## Author contributions

R.A. designed and performed the experiments, analyzed data, and wrote the manuscript. A.T. generated biotin-PEG-functionalized coverslips and adapted the initial conditions to visualize tubulin polymerization on these surfaces. S.P. contributed to research design, mentoring and wrote the manuscript.

## Ethics

Animal experimentation: This study was performed in strict accordance with the recommendations in the Guide for the Care and Use of Laboratory Animals of the National Institutes of Health. All of the animals were handled according to approved Institutional Animal Care and Use Committee (IACUC) protocol # 1941-16 of Princeton University.

## Figure Supplement Legends

**Figure 1 – figure supplement 1.**
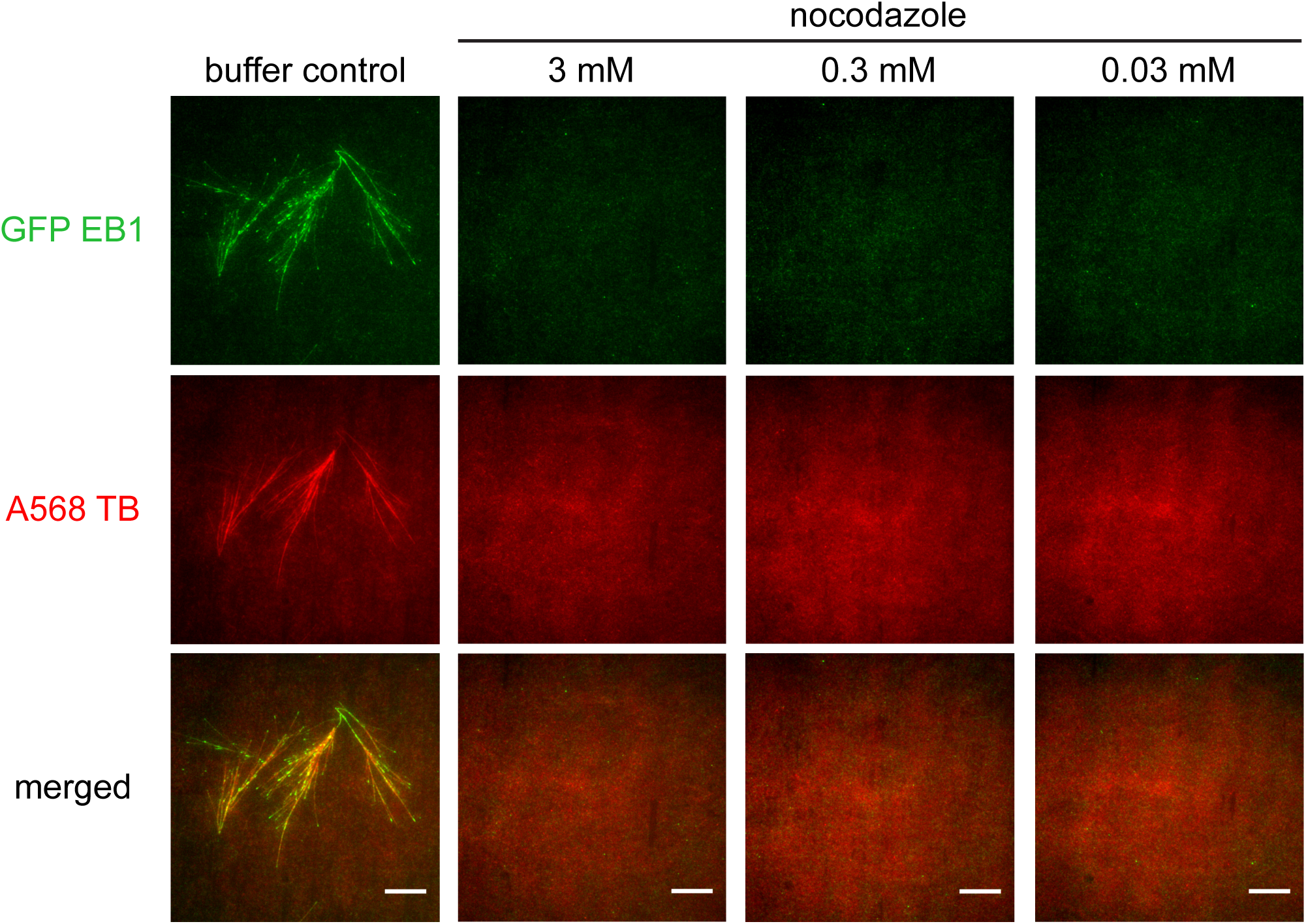
Testing the inhibitory effect of nocodazole in Xenopus egg extract. Branching microtubule nucleation was stimulated in Xenopus egg extract with 10 μM RanQ69L in the presence of increasing concentrations of nocodazole. microtubules were labeled with Alexa568 tubulin (red) and their plus-ends with EB1-GFP (green). Scale bars, 10 μm. The experiment was repeated three times with different Xenopus egg extracts

**Figure 1 – figure supplement 2.**
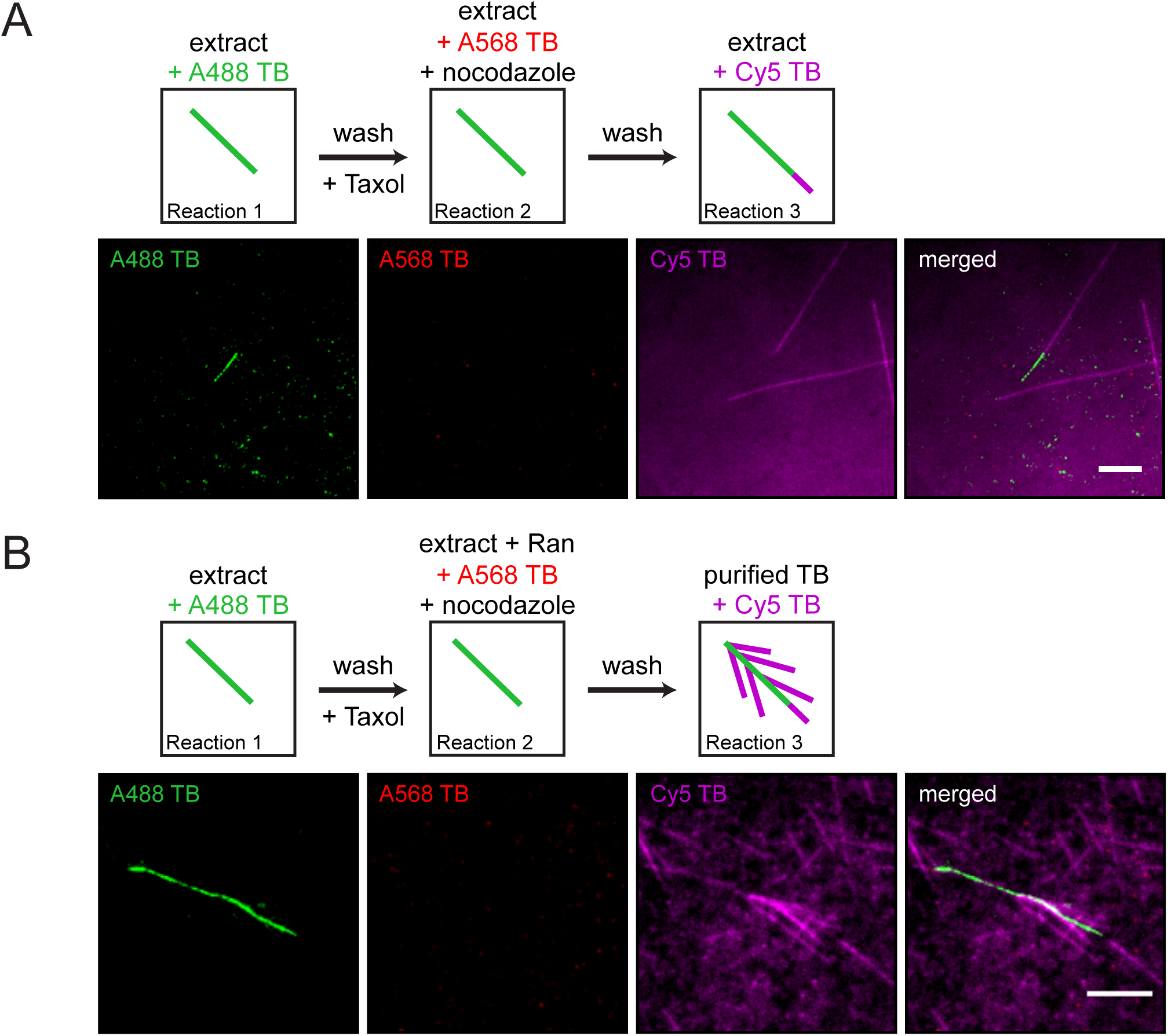
Sequential Xenopus egg extract reactions. (**A**) Single microtubules formed on the glass surface in the first extract supplemented with Alexa488 tubulin (green). A second extract supplemented with Alexa568 tubulin (red) and nocodazole, but lacking RanQ69L, was subsequently introduced, followed by a third extract supplemented with Cy5 tubulin (magenta). Pre-existing microtubules (green) only extended from their plus-ends (magenta) in the third extract reaction because no branching factors were released in the second reaction step, while no microtubules formed in the presence of nocodazole (red). (**B**) Analogous to (A), except that the second extract was supplemented with RanQ69L, and the third extract reaction was substituted for purified Cy5 tubulin (magenta). Branched microtubules (magenta) nucleated from pre-existing microtubules (green), while no microtubules formed in the presence of nocodazole (red). For all experiments, images were collected approximately 5 min after the last solution was exchanged. Scale bars, 5 μm. The experiments were repeated three times with different Xenopus egg extracts.

**Figure 2 – figure supplement 1.**
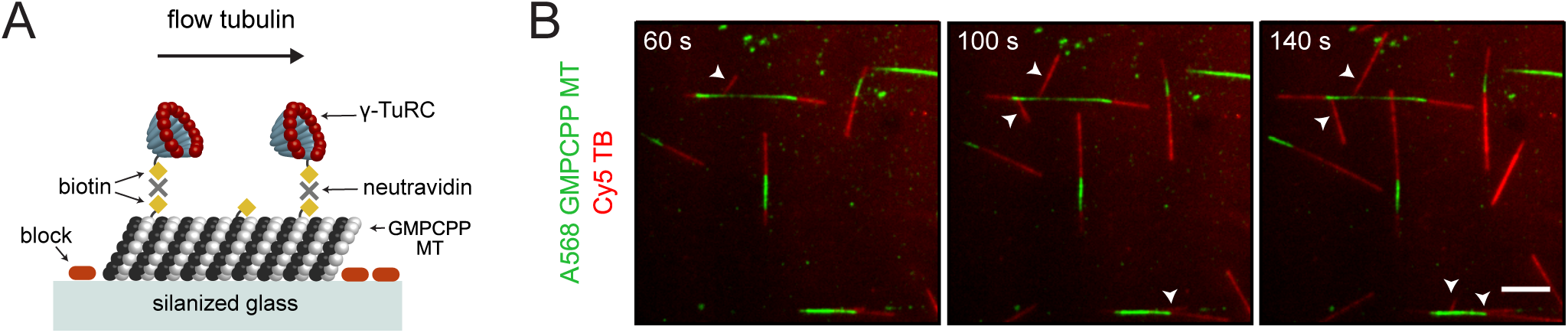
Microtubule nucleation from artificially-attached γ-TuRCs to a template microtubule. (**A**) Diagram of the experimental set-up. GMPCPP-stabilized microtubules attached non-specifically to a silanized cover glass, and γ-TuRCs attached to the microtubules with biotin-neutravidin links. Nucleation of new microtubules was visualized using Cy5 tubulin. (**B**) Using the set-up in (A), the formation of artificial microtubule branches (red, arrowheads) from GMPCPP-stabilized microtubules (green) was observed. Scale bar, 5 μm. Experiment was performed once.

**Figure 2 – figure supplement 2.**
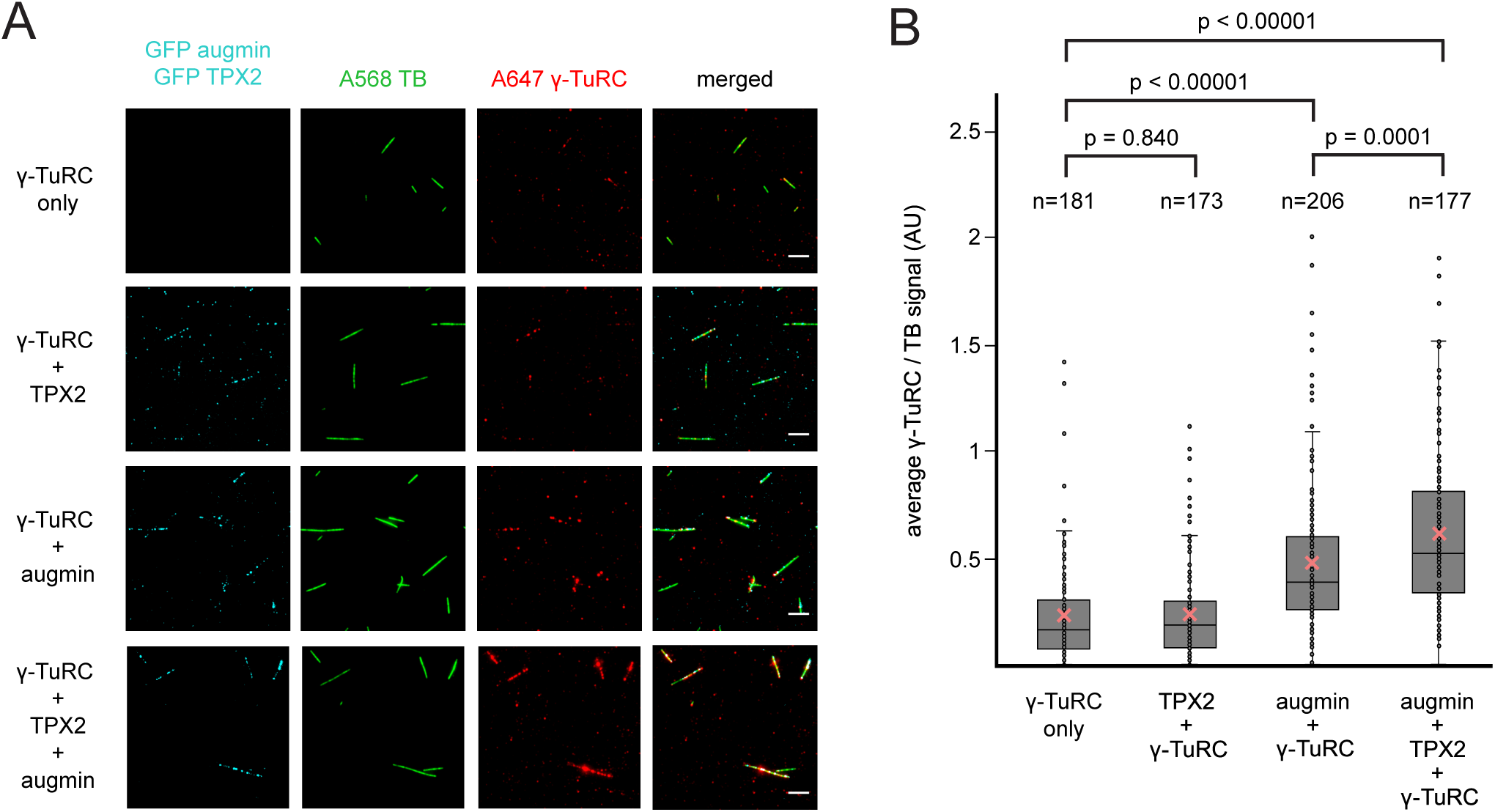
Recruitment of γ-TuRC to a template microtubule by augmin and TPX2. (**A**) γ-TuRC visualized using Alexa647-labeled antibodies (red) along microtubules (green), in the absence or presence of GFP-augmin and GFP-TPX2 (cyan). Scale bars, 5 μm. (**B**) Boxplot of average γ-TuRC signal relative to the average tubulin signal, where each dot represents one microtubule from the experiment in (A). The number of microtubules (n) was obtained from one experiment. the boxes extend from 25th to 75th percentiles, the whiskers extend from minimum to maximum values, and the mean values are plotted as crosses. P-values were calculated from independent T-tests.

**Figure 3 – figure supplement 1.**
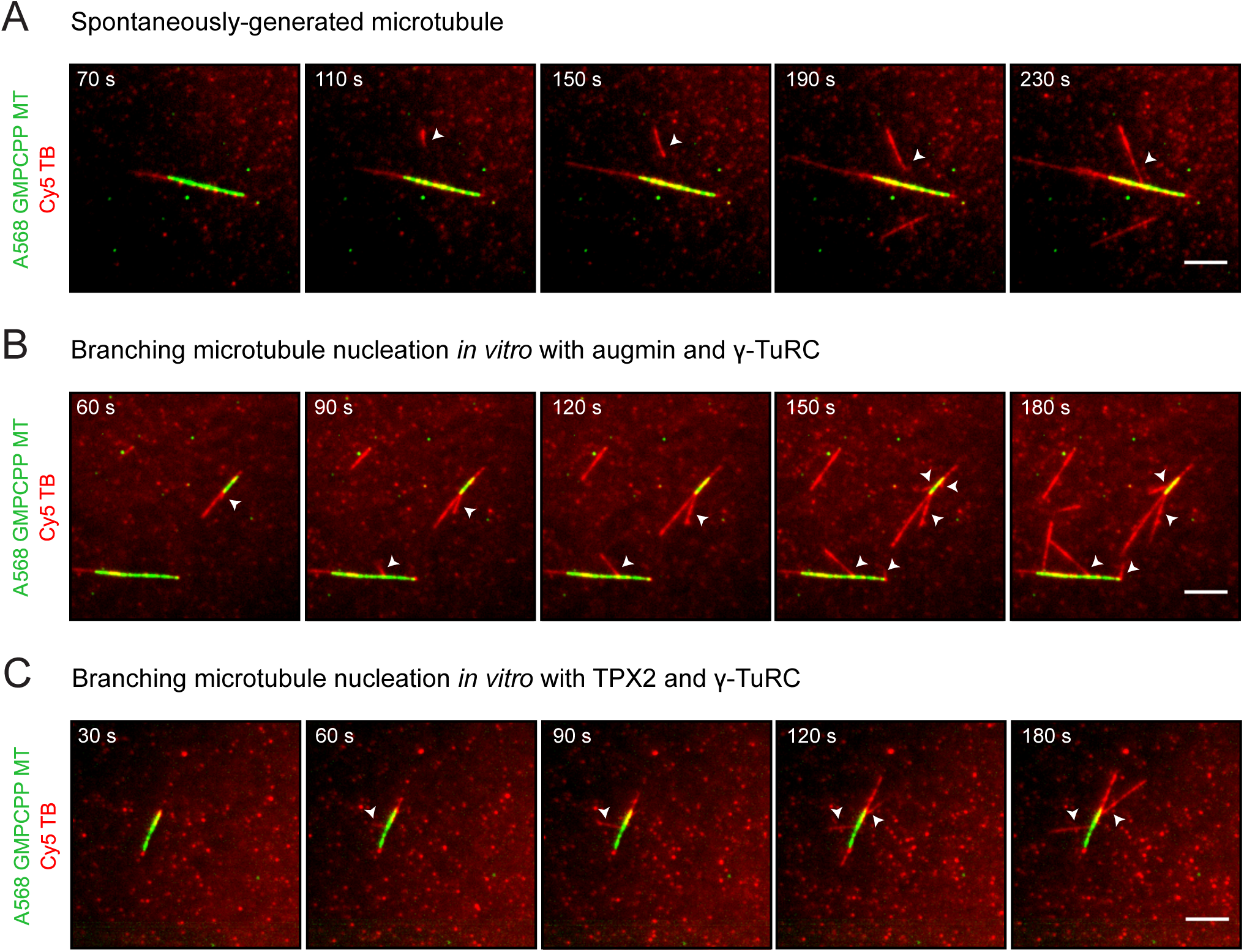
Microtubules can spontaneously form in solution and subsequently interact with the template GMPCPP-stabilized microtubule. (**A**) Time-lapse images from the experiment in Fig. 3B showing an example of a microtubule (red, arrowhead) that is spontaneously nucleated in solution and contacts the GMPCPP-stabilized template microtubule (green) afterwards. (**B**) Similar to the experiment in Fig. 3B, but only augmin and γ-TuRC were bound to the GMPCPP-stabilized microtubule. The formation of some microtubule branches (red, arrowheads) from GMPCPP-stabilized microtubules (green) was observed. (**C**) Similar to the experiment in Fig. 3B, but only TPX2 and γ-TuRC were bound to the GMPCPP-stabilized microtubule. The formation of some microtubule branches (red, arrowheads) from GMPCPP-stabilized microtubules (green) was also observed. Scale bars, 5 μm.

**Figure 3 – figure supplement 2.**
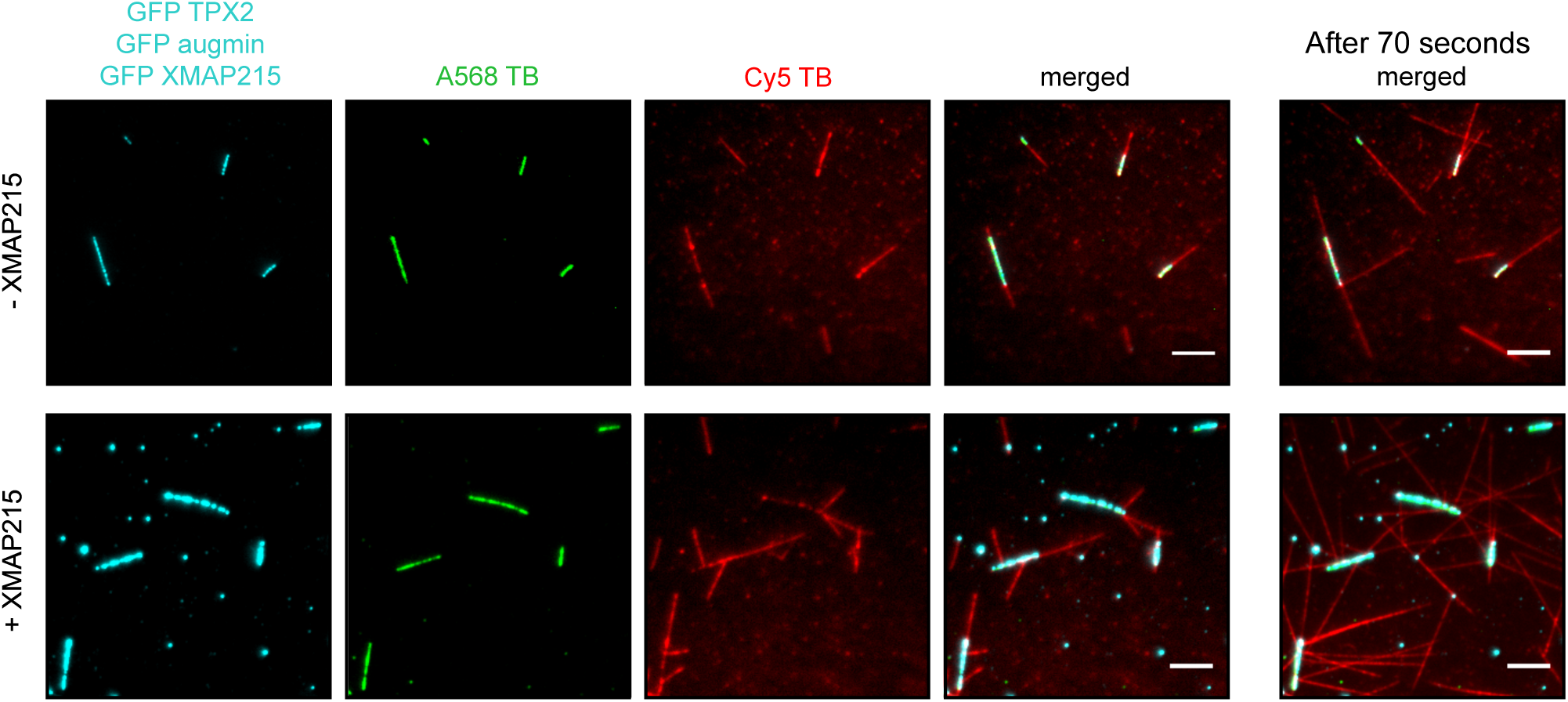
Reconstitution of branching microtubule nucleation using purified augmin, TPX2, γ-TuRC and XMAP215. Similar to the experiment in Fig. 3B, but comparing the effect of having GFP-XMAP215 in the final solution of Cy5 tubulin. The images correspond to the first frame of the time-lapse collected. The panels on the right (merged only) correspond to the same fields of view 70 seconds later. Scale bar, 5 μm. Experiment was performed once.

## Video Legends

**Video 1. Branching microtubule nucleation from a pre-existing microtubule in Xenopus egg extract (related to Figure 1A).** A single microtubule formed on the glass surface in extract supplemented with Alexa488 tubulin (green). A second extract supplemented with Alexa568 tubulin (red) and RanQ69L was subsequently introduced. Branched microtubules (red) nucleated from the pre-existing microtubule (green). The sample was imaged every 2 sec. Scale bar, 10 μm.

**Video 2. The proteins necessary for branching microtubule nucleation in Xenopus egg extract bind to a pre-existing microtubule preceding and independent of the nucleation event (related to Figure 1C).** Single microtubules formed on the glass surface in extract supplemented with Alexa568 tubulin (green). A second extract supplemented with RanQ69L and nocodazole was subsequently introduced during which branching factors bound to the pre-existing microtubule. Finally, a mixture of purified Cy5 tubulin (red) and XMAP215 was added. Branched microtubules (red) nucleated from pre-existing microtubules (green). The sample was imaged every 2 sec. Scale bar, 10 μm.

**Video 3. Reconstitution of branching microtubule nucleation using purified augmin, TPX2 and γ-TuRC (related to Figure 3B).** A GMPCPP-stabilized microtubule (green) with bound augmin, TPX2, and γ-TuRC, served as a template for the nucleation of branched microtubules (red). The sample was imaged every 2 sec. Scale bar, 5 μm.

